# Synergistic activities of ethanolic extracts of *Jatropha tanjorensis* leaves in conventional management of rheumatoid arthritis in the ankles of Wistar Rats

**DOI:** 10.1101/2022.07.22.501197

**Authors:** Chiadikobi Lawrence Ozoemena, Ifeanyi Anthony Egwuatu, Ifeanacho Ezeteonu Abireh, Elizabeth Finbarrs-Bello, Ignatius Ikemefuna Ozor, Anthony Okechukwu Akpa

## Abstract

**Background:** Rheumatoid arthritis (RA) is a common cause of chronic inflammatory joint disease. Plant extracts contain several bioactive factors which can re-establish the homeostasis of joints and joint cartilages.

**Objectives:** This study aim to evaluate synergism of ethanolic extracts of *Jatropha tanjorensis* leaves with Ibuprofen and Sulfasalazine as an antirheumatic agent.

**Methodology:** 25 adult male Wistar rats were subjected to different types of treatment (110 days); A: Positive control, B: Ibuprofen, Sulfasalazine and Low dose of extract, C: Ibuprofen, Sulfasalazine and Medium dose of extract, D: Ibuprofen, Sulfasalazine and High dose of extract, E: Normal control. Groups A-D were collagen-induced arthritic (CIA) models. One week before sacrifice, anterior-posterior and lateral diameters of both ankles, physical appearance and weight were assessed. After sacrifice, histological analysis of ankles using modified Mankin scoring system was done.

**Results:** Groups B, C and D had significant improvements in direct proportion to the dosages of J. tanjorensis extract administered. Increased doses slowed down the progression of cartilage destruction evidenced by prevention of joint swellings and preservation of chondrocytes and its histological features. Group A (positive control) exhibited cartilage destruction but no cartilage changes noted in group E (normal control).

**Conclusion:** We demonstrated synergistic effects of J. tanjorensis with Ibuprofen and Sulfasalazine combination therapy on animal model of collagen-induced rheumatoid arthritis. The evaluation of these effects were assessed by several means; physical appearance, assessment of weight, anterior-posterior and lateral diameters, and histological examination of both ankles of the animal that characterizes the originality of the study.

**Summary:** What is already known on this topic – The current therapeutic regimen for RA has some disadvantageous side effects.

What this study adds – An alternative to the conventional management of RA with lower side effects.

How this study might affect research, practice or policy – There is need to standardize dosages of newer herbs identified to be therapeutic.

## INTRODUCTION

The cultivation of herbs has been ascribed to the discovery of plants rich in pharmacologically active constituents with an edge that is proven against well-known disease entities and medical conditions [1]. Hundreds of species of tropical angiosperms grown have medicinal properties, thereby making traditional medicine more available to the general populace in comparison to modern medicine [2]. The use of traditional herbs in the treatment of diseases is largely due to their nutraceutical components which is attributed to plant secondary metabolites [3, 4]. In Nigeria, some claims have been laid on the efficiency of *Jatropha tanjorensis* in ameliorating oxidative stress [3]; a perennial herb that belongs to the Euphorbiaceae family. It is commonly used in Nigeria as a blood building vegetable, anti-plasmodial and anti-oxidant against oxidative stress [5, 6, 7]. Granted that many studies have been pursued to a conclusion on *Jatropha tanjorensis*, howbeit, none of these numerous studies addressed the therapeutic potentials of ethanolic extract of *Jatropha tanjorensis* leaves on autoimmune diseases.

Rheumatoid arthritis (RA) is the most common cause of chronic inflammatory joint disease; an autoimmune systemic disease with slow but progressive destruction of joints and joint cartilages [8]. There is no known cure to RA presently, thus the adoption of supportive and palliative approaches in its management with an aim to slow down disease progression, alleviate symptoms and reduce functional limitations [9].

In view of the anabolic actions and synergistic effects of therapeutic options on damaged cartilage from RA, ethanolic extract of *Jatropha tanjorensis* is a compelling candidate to investigate as a therapeutic option, because it contains several bioactive factors from its secondary metabolites which can reestablish the homeostasis of the synovial fluid [10]. It is also a rich source of antioxidant nutrients like phosphorus, selenium, zinc, vitamin C and vitamin E; these micronutrients assists in inflammation modulation and cell migration induction [9]. Its Phytochemical components contain secondary metabolites including but not limited to; phytates, phenols, alkaloids, terpenoids, saponins, flavonoids and tannins which contributes to the inhibition of progressive damage of joint capsules in the management of rheumatoid arthritis disease [11-14]. Flavanoids are attractive because of their anti-inflammatory, anti-microbial and anti-oxidant potentials as secondary metabolites [12, 15, 16]. Also, with the prevalence of RA mostly involving geriatrics, immunosuppressants such as infliximab and its class of drugs is not totally the best of option in the approach to combat rheumatoid arthritis due to their side effects [17].

In this study, we used the Collage-induced arthritis (CIA) model which represents the gold standard RA animal model because its pathological features closely resemble those in human; synovium lymphocyte infiltration and hyperplasia with cell swelling [18].

We report in this study with intent to evaluate the use of *Jatropha tanjorensis* in the management of RA for these aforementioned reasons; being cost-effective, readily available and ease of preparation. We evaluated the potentials of ameliorating the side effects of both Non-steroidal Anti-Inflammatory Drugs (NSAIDs) and Disease Modifying Anti-Rheumatic Drugs (DMARDs) which are currently used in the management of RA while enhancing therapeutic efficacy using *J. tanjorensis*. The evaluation concerned mainly; physical appearance, weight, anthropometry which included measuring the anterior-posterior and lateral diameters of both ankles, and histological examination of both ankle joints of the animals.

## MATERIALS AND METHODOLOGY

### Plant Material and Extract

*Jatropha tanjorensis* leaves were obtained from a farm settlement in Mbanugo, Coal Camp, Enugu state of Nigeria, between May 2021 and October, 2021. The air dried and ground leaves (2500g) was cold-macerated and extracted in 6 volumes of absolute alcohol to give the extract which was concentrated in a hot water bath at 80°c (mean yield: 38.26±0.89g w/w), and stored in a sample bottle at -4°c until ready for use. Phytochemical analysis gave positive tests for phytates, phenols, terpenoids, saponins, flavonoids and tannins.

### Experimental Animals and treatment

A total of 50 adult male Wistar rats weighing 180-250g were housed in cross-ventilated cages in the animal house of Department of Anatomy, Enugu State University of Science and Technology, Enugu. Conditions of the facility were 20-28°c with 12 hour light/dark cycle. Animals had access to commercial growers mash and tap water ad libitum. All animals were fed and acclimatized for three weeks before the model construction. 50 rats were grouped accordingly;

1. 5 rats - Pilot dose-response trial of Bovine CII-CFA emulsion by injection at the base of the tail before determining appropriate dose due to variations in dose requirements in various strains of rodents.
2. 5 rats - Normal control group.
3. 40 rats - Model construction.

The rats were anaesthetized using 5% isoflurane in an induction-bell jar. For model construction, 200µl of bovine CII-CFA emulsion (0.2mg/ml) was injected into the base of the tail using a 26F-guage needle. Booster dosage was performed 9 days later by injecting 100µl of bovine CII-CFA emulsion (0.1mg/ml). Rats in the normal control group received equal volume of normal saline.

### Evaluating rheumatoid arthritis in the rats

On day 1 and day 16 (1 week after booster dose was given), the rats physical appearance was assessed and their limbs were measured using a vernier caliper to check for possible limb swellings. Arthritis index (AI) was determined to be positive and models considered successful when limb swellings involved an entire limb, affecting multiple joints symmetrically. The anterior-posterior diameter (between the posterior strip of the ankle and heel) and lateral diameter of the ankle of both hindlimbs were measured on days 0 and 16. Also on day 16, blood samples from 5 rats chosen randomly from the model group were investigated for anti-cyclic citrullinated peptide antibody (ACCPA). They were all positive with average value of 49 units.

### Administration of *J. tanjorensis* extract, Ibuprofen and Sulfasalazine

At day 16, 20 successful models were selected from the model group and randomized into four treatment groups (A-D) with five animals each. All groups were subjected to different types of treatment by oro-gastric intubation for 12 weeks as follows:

### Positive control (Group A) – distilled water only

Low dose (Group B) – Ibuprofen (5mg/kg/day), Sulfasalazine (5mg/kg/day), and *J. tanjorensis* extract at a dose of 0.5g/kg/day corresponding to 1ml/kg body weight.

Medium dose (Group C) – Ibuprofen (5mg/kg/day), Sulfasalazine (5mg/kg/day), and *J. tanjorensis* extract at a dose of 1g/kg/day corresponding to 2ml/kg body weight.

High dose (Group D) – Ibuprofen (5mg/kg/day), Sulfasalazine (5mg/kg/day), and *J. tanjorensis* extract at a dose of 1.5g/kg/day corresponding to 3ml/kg body weight.

Normal control (Group E) – distilled water only.

At day 93, the anterior-posterior and lateral diameters of both hindlimbs were measured again. At day 100, all rats were sacrificed by cervical dislocation and both ankles rapidly dissected and processed for histological study.

### Histological study

Tissues of the ankles were fixed in 10% buffered formalin and post-fixed in zenker’s fluid for 18 hours, and then processed for paraffin sectioning. Sections were cut at 5µ in thickness and stained using Hematoxylin and Eosin (H&E) dyes. The slides were analyzed and scored using the Modified Mankin scoring system.

## RESULTS

### General observations (day 0 to day 16)

The observation in the physical appearance of the model group on day 0 were yellowish patches of fur colour changes (2cm by 2cm). Some rats had ulceration and scab formation at the site of injection measuring about 0.5cm by 1 cm on average. Day 16 marked an increase in the size of the yellowish discolouration of furs to about 4cm by 5cm. Their joints were also swollen and they exhibited limited mobility.

### Changes observed in body weight

At day 0, the body weight between the rats in all groups had no significant difference (P > 0.05). However, by day 16, the body weights of the rats in the model group had a significant decrease in comparison to the normal control group (P ≤ 0.05). By day 93, the body weights of rats in groups A and B were significantly decreased in comparison to other groups. While groups C and D showed significant improvement in weight and were significantly different in comparison to group A (P ≤ 0.05) (Table 1).

**Table 1:**
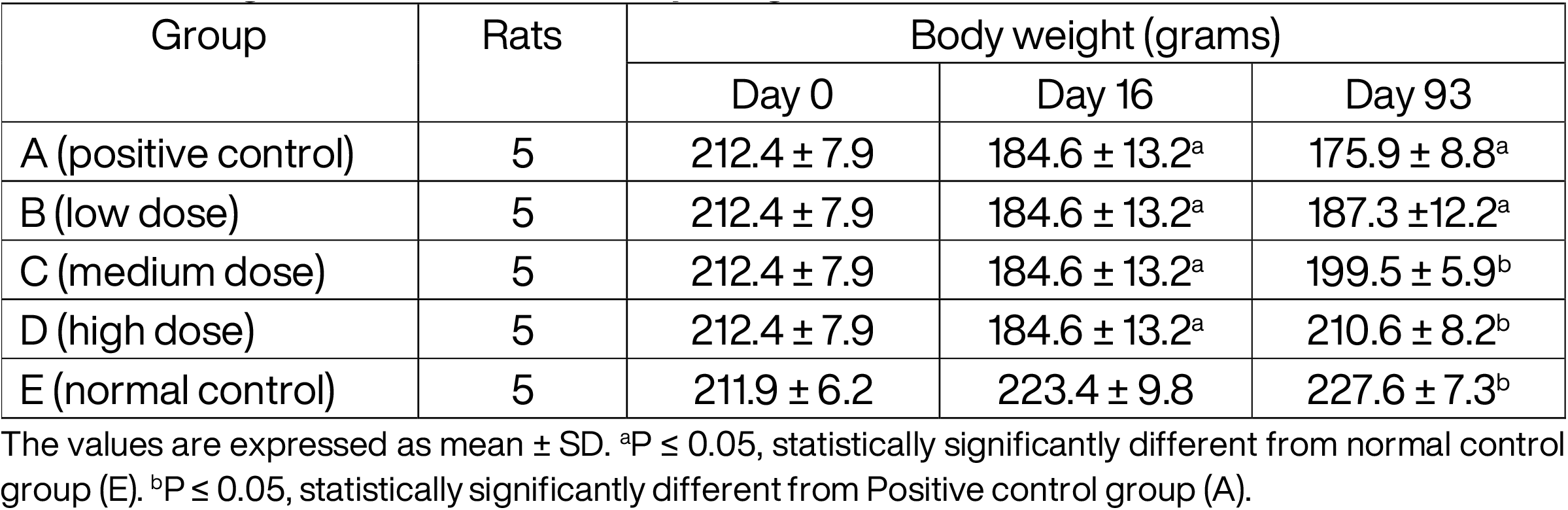
Changes observed in the body weight of rats

| Group | Rats | Body weight (grams) |  |  |
| --- | --- | --- | --- | --- |
|  |  | Day 0 | Day 16 | Day 93 |
| A (positive control) | 5 | 212.4 ± 7.9 | 184.6 ± 13.2<sup>a</sup> | 175.9 ± 8.8<sup>a</sup> |
| B (low dose) | 5 | 212.4 ± 7.9 | 184.6 ± 13.2<sup>a</sup> | 187.3 ± 12.2<sup>a</sup> |
| C (medium dose) | 5 | 212.4 ± 7.9 | 184.6 ± 13.2<sup>a</sup> | 199.5 ± 5.9<sup>b</sup> |
| D (high dose) | 5 | 212.4 ± 7.9 | 184.6 ± 13.2<sup>a</sup> | 210.6 ± 8.2<sup>b</sup> |
| E (normal control) | 5 | 211.9 ± 6.2 | 223.4 ± 9.8 | 227.6 ± 7.3<sup>b</sup> |

The values are expressed as mean ± SD. <sup>a</sup>P ≤ 0.05, statistically significantly different from normal control group (E). <sup>b</sup>P ≤ 0.05, statistically significantly different from Positive control group (A).

### Changes observed in swelling of limbs in the treatment groups

At day 0, there was no significant difference in the anterior-posterior and lateral diameters of both ankles of the rats in all groups (P > 0.05). However, by day 16, the anterior-posterior and lateral diameters of the ankles of the hind-limbs in the model groups were significantly larger than that of the normal control group (P ≤ 0.05). By day 93, the anterior-posterior and lateral diameters of both ankles measured in groups A and B were significantly larger than the rest of the groups (P ≤ 0.05). Group D was significantly different when compared to the positive control group (A). However, Group C was significantly different in comparison to both group A and E (positive and normal control groups) (P ≤ 0.05) (Table 2).

**Table 2:** Changes observed in swelling of the ankles of the hind-limbs of rats

| Ankles | Group | n | Anterior-posterior diameter (mm) |  |  | Lateral diameter (mm) |  |  |
| --- | --- | --- | --- | --- | --- | --- | --- | --- |
|  |  |  | Day 0 | Day 16 | Day 93 | Day 0 | Day 16 | Day 93 |
| Right ankles | A (positive control) | 5 | 7.54 ± 0.65 | 11.94 ± 0.38<sup>a</sup> | 14.23 ± 0.41<sup>a</sup> | 6.32 ± 0.56 | 8.92 ± 0.46<sup>a</sup> | 13.89 ± 0.71<sup>a</sup> |
|  | B (low dose) | 5 | 7.54 ± 0.65 | 11.94 ± 0.38<sup>a</sup> | 12.18 ± 0.67<sup>a</sup> | 6.32 ± 0.56 | 8.92 ± 0.46<sup>a</sup> | 11.91 ± 0.74<sup>a</sup> |
|  | C (medium dose) | 5 | 7.54 ± 0.65 | 11.94 ± 0.38<sup>a</sup> | 9.72 ± 0.61<sup>a,b</sup> | 6.32 ± 0.56 | 8.92 ± 0.46<sup>a</sup> | 7.91 ± 0.32<sup>a,b</sup> |
|  | D (high dose) | 5 | 7.54 ± 0.65 | 11.94 ± 0.38<sup>a</sup> | 8.21 ± 0.85<sup>b</sup> | 6.32 ± 0.56 | 8.92 ± 0.46<sup>a</sup> | 7.1 ± 0.68<sup>b</sup> |
|  | E (normal control) | 5 | 7.49 ± 0.43 | 7.66 ± 0.29 | 7.73 ± 0.46<sup>b</sup> | 6.42 ± 0.42 | 6.62 ± 0.77 | 6.54 ± 0.49<sup>b</sup> |
| Left ankles | A (positive control) | 5 | 7.62 ± 0.57 | 12.05 ± 0.71<sup>a</sup> | 13.97 ± 0.11<sup>a</sup> | 6.29 ± 0.74 | 9.19 ± 0.54<sup>a</sup> | 13.71 ± 0.48<sup>a</sup> |
|  | B (low dose) | 5 | 7.62 ± 0.57 | 12.05 ± 0.71<sup>a</sup> | 12.43 ± 0.71<sup>a</sup> | 6.29 ± 0.74 | 9.19 ± 0.54<sup>a</sup> | 11.76 ± 0.81<sup>a</sup> |
|  | C (medium dose) | 5 | 7.62 ± 0.57 | 12.05 ± 0.71<sup>a</sup> | 9.61 ± 0.32<sup>a,b</sup> | 6.29 ± 0.74 | 9.19 ± 0.54<sup>a</sup> | 7.69 ± 0.17<sup>a,b</sup> |
|  | D (high dose) | 5 | 7.62 ± 0.57 | 12.05 ± 0.71<sup>a</sup> | 7.91 ± 0.83<sup>b</sup> | 6.29 ± 0.74 | 9.19 ± 0.54<sup>a</sup> | 6.82 ± 0.31<sup>b</sup> |
|  | E (normal control) | 5 | 7.58 ± 0.66 | 7.75 ± 0.36 | 7.64 ± 0.52<sup>b</sup> | 6.32 ± 0.43 | 6.58 ± 0.27 | 6.45 ± 0.36<sup>b</sup> |

The values are expressed as mean ± SD. <sup>a</sup>P ≤ 0.05, statistically significantly different from normal control group (E). <sup>b</sup>P ≤ 0.05, statistically significantly different from Positive control group (A). <sup>a,b</sup>P ≤ 0.05, statistically significantly different from Positive control group (A) and Normal control group (E).

### Light microscopic analysis with modified Mankin scoring system

Paraffin sections of the knee joints of ethanolic *J. tanjorensis* leaf extract-treated rats showed that the extract produced a significant dose-dependent improvement in the cartilage erosion, peripheral staining of chondrocytes, spatial arrangement of chondrocytes and background staining intensity of the matrix. Micrographs from group A had features suggestive of massive cartilage destruction which included erosion of calcified cartilages, intensely enhanced chondrocyte periphery, sparsely distributed chondrocytes, no background staining; giving a total score of 14/15 (figure 1). With an extract dose of 0.5g/kgbw, the extent of cartilage destruction in group B was limited to a Mankin Score of 8/15 with superficial fibrillation of cartilages noted, intensely enhanced chondrocyte periphery staining, clustered spatial arrangement of chondrocytes and moderately reduced background staining intensity (figure 2). Doubling the dosage of the extract (1.0g/kgbw) showed a remarkable decrease in the extent of cartilage damage I group C with a Mankin Score of 5/15. The cartilages showed superficial fibrillation, however the chondrocyte periphery staining was slightly enhanced with diffuse hyper-cellular spatial arrangement of the chondrocytes and slightly reduced background staining intensity (figure 3). At a dosage of 1.5g/kgbw the Mankin Score improved to 3/15 in group D. The cartilages appeared rough but non-eroded with a normal chondrocyte periphery staining, diffuse hyper-cellular spatial arrangement of chondrocytes and slightly reduced background staining intensity (figure 4). There was no remarkable feature noted in the micrograph of group E (normal control) (figure 5).

**Figure 1:**
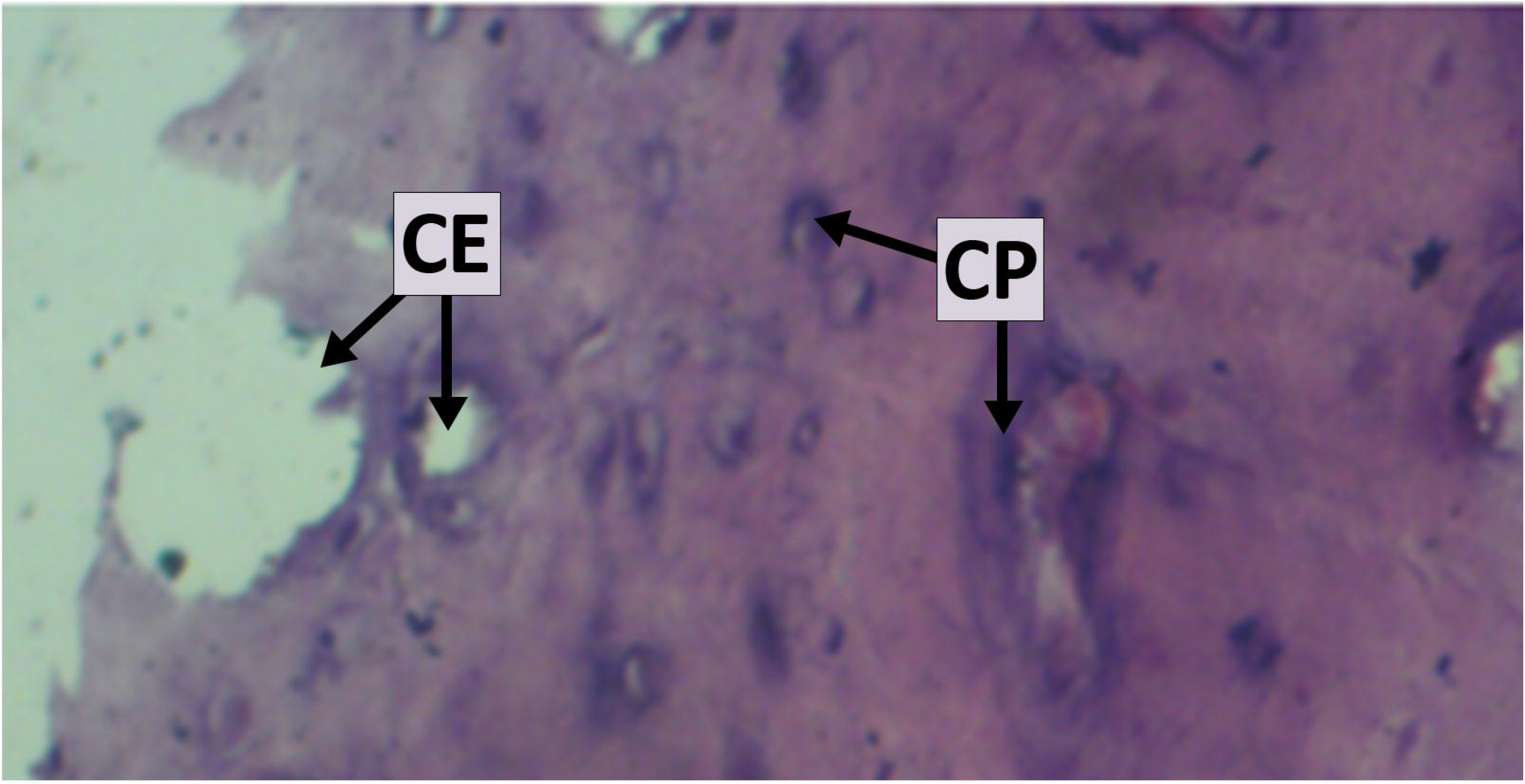
Micrograph of joint cartilage in group A (positive control) showing features suggestive of massive cartilage destruction. CE = cartilage erosion extending into calcified cartilage, with clustering chondrocytes spatial arrangement. CP = intensely enhanced chondrocyte periphery staining. Background intensity staining showing no dye absorption. [H & E Stain x 400].

**Figure 2:**
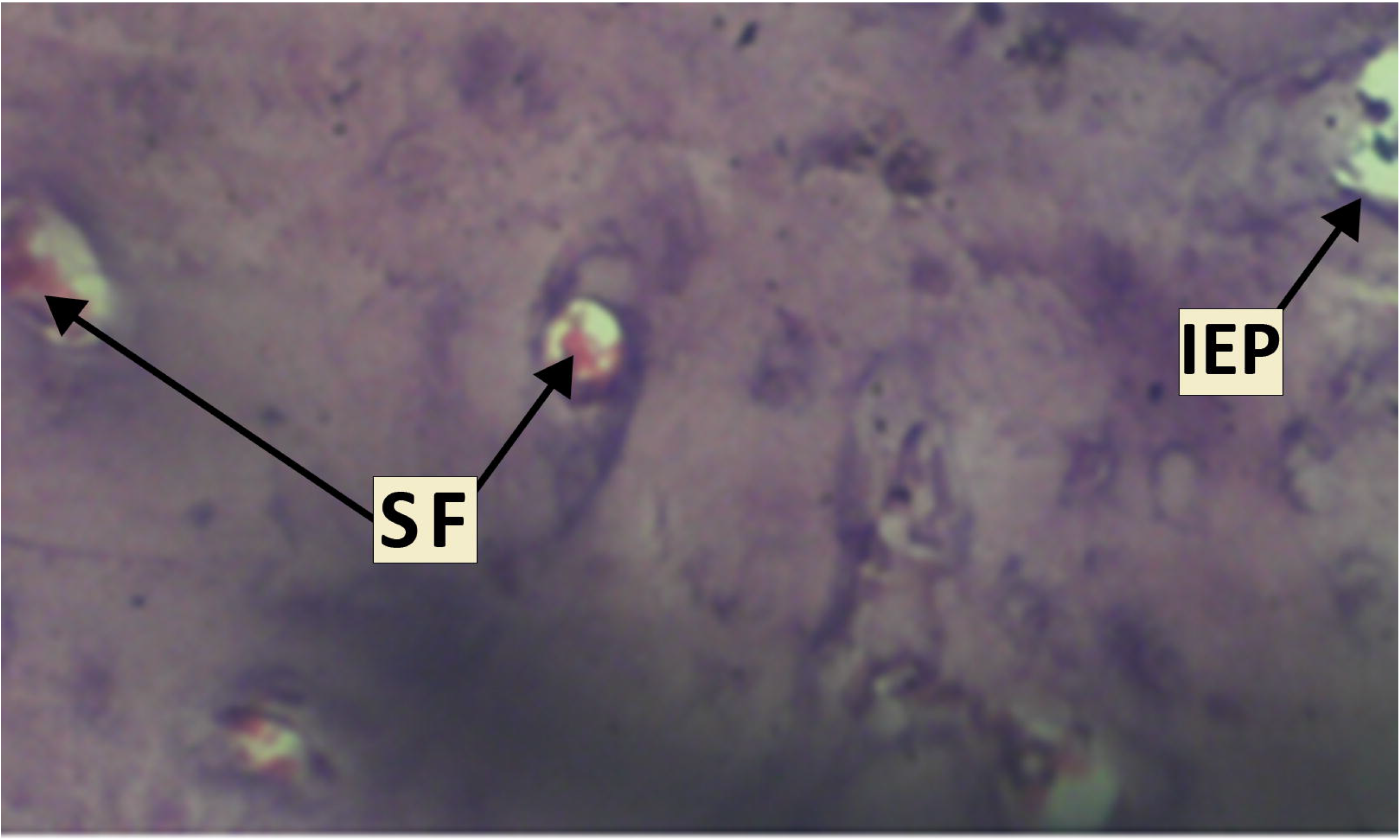
Micrograph of joint cartilage in group B (0.5g/kg of *J. tanjorensis* leaf extract) showing cartilage destruction; SF = superficial fibrillation of cartilage, IEP = intensely enhanced chondrocyte periphery staining, clustering arrangement of chondrocytes and moderately enhanced background staining intensity [H & E Stain x 400].

**Figure 3:**
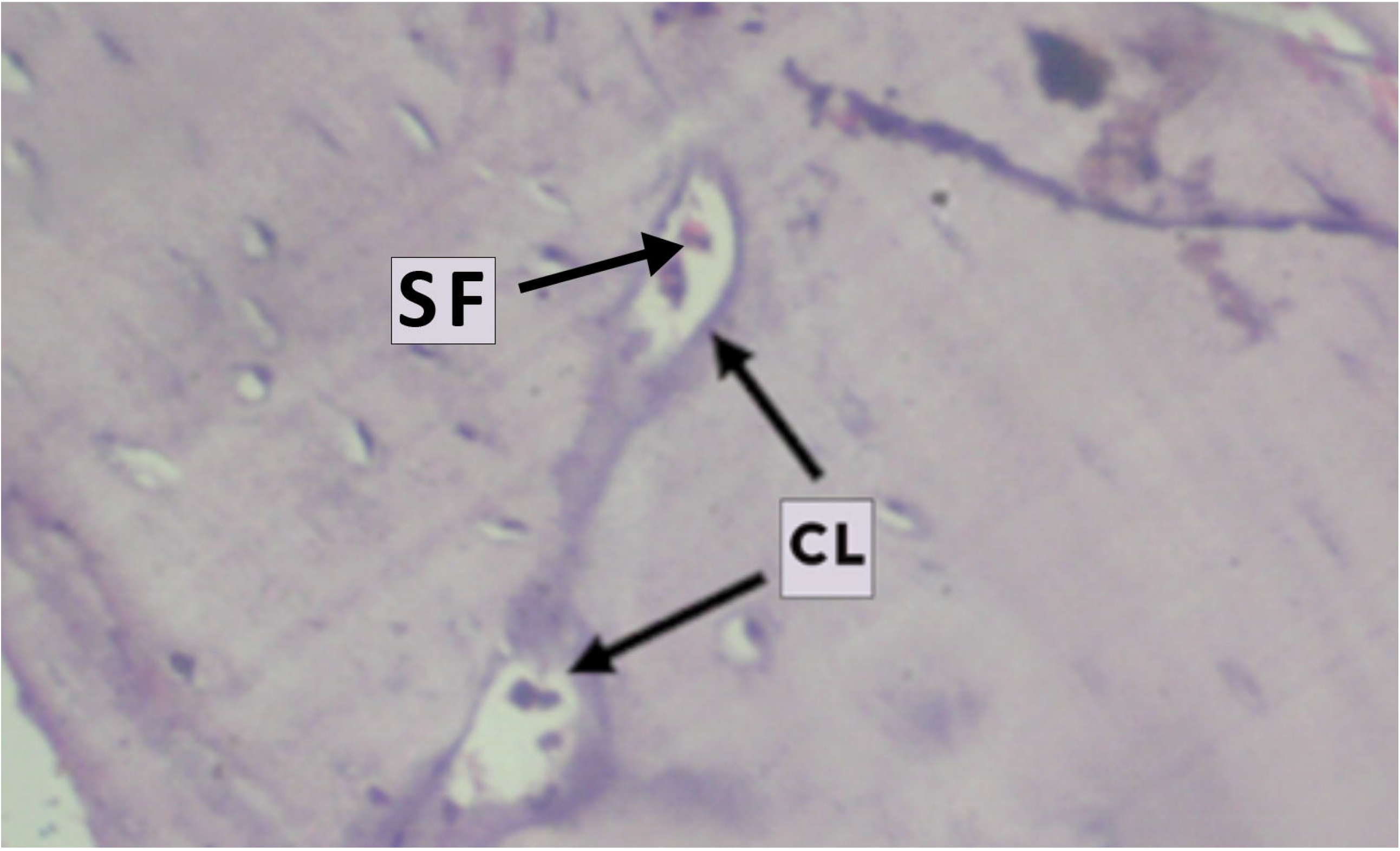
Micrograph of joint cartilage in group C (1.0g/kg of *J. tanjorensis* leaf extract) showing moderate cartilage destruction; SF = superficial fibrillation of cartilage, CL = chondrocyte lacunae with slightly enhanced periphery staining, diffuse hyper-cellular spatial arrangement of chondrocytes and slightly reduced background staining intensity [H & E Stain x 400].

**Figure 4:**
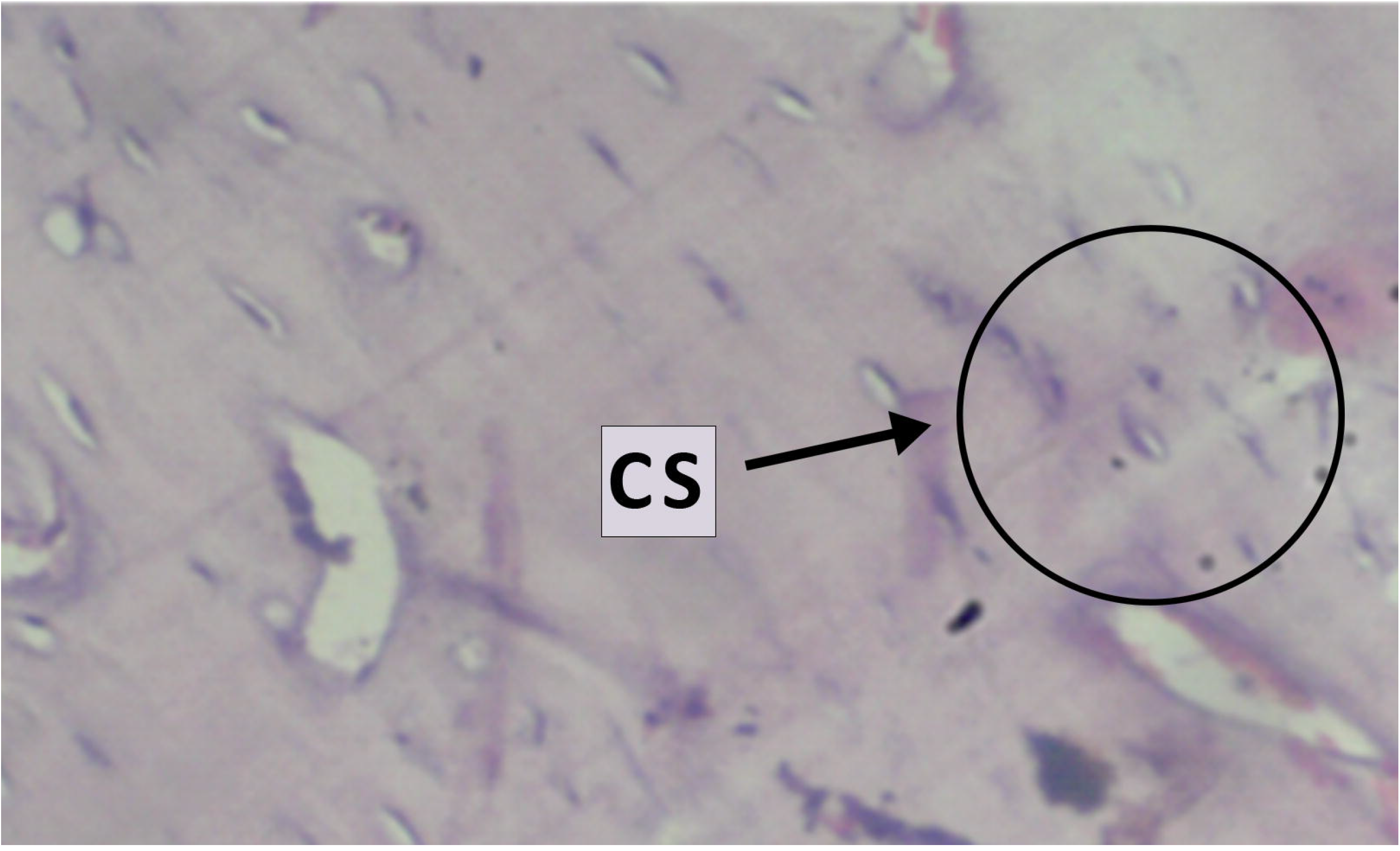
Micrograph of joint cartilage in group D (1.5g/kg of *J. tanjorensis* leaf extract) showing no cartilage destruction; rough non-eroded cartilages, normal chondrocyte periphery staining, CS = diffuse hyper-cellular spatial arrangement of chondrocytes and slightly reduced background staining intensity [H & E Stain x 400].

**Figure 5:**
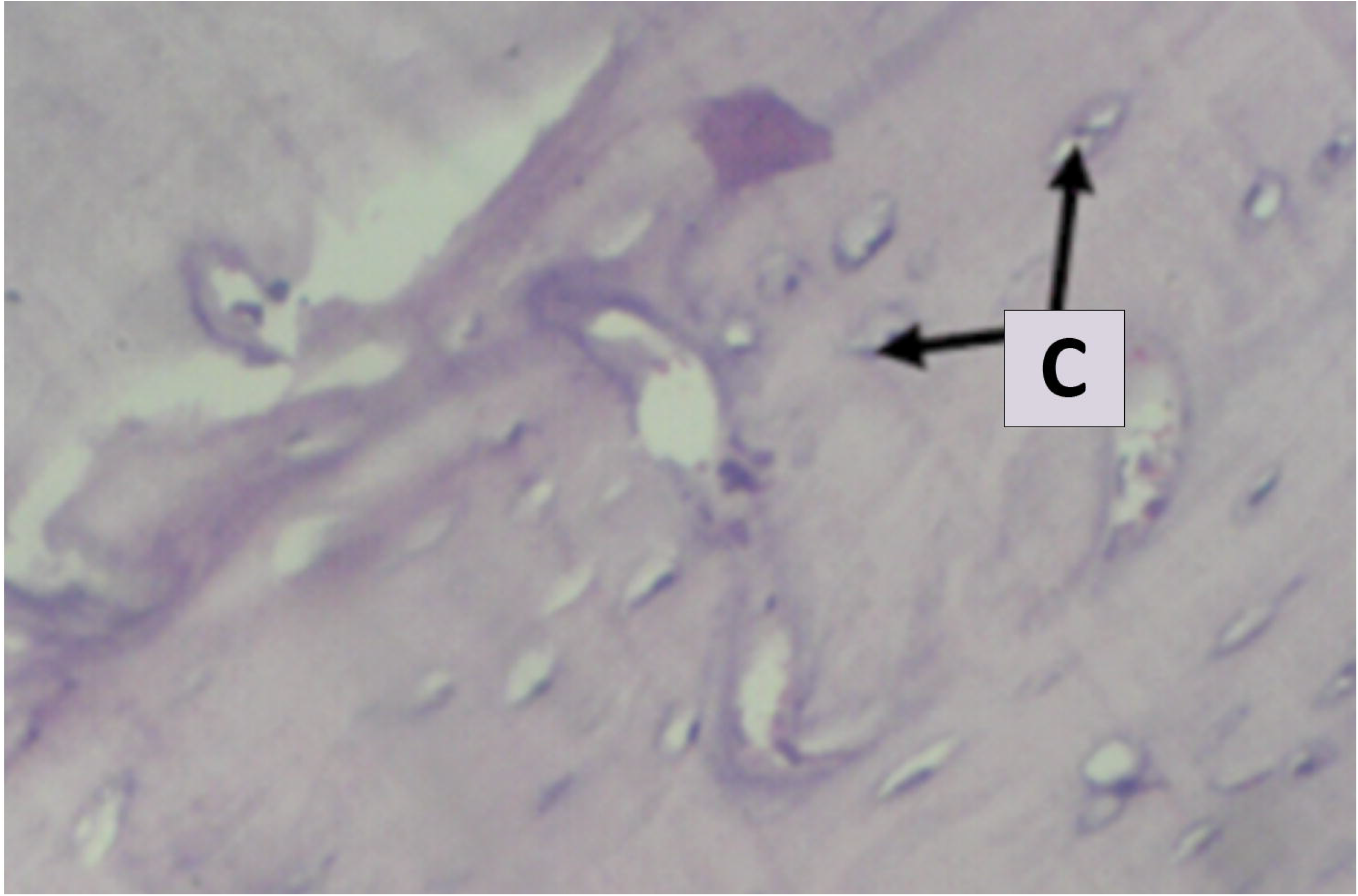
Micrograph of joint cartilage in group E (normal control) showing no remarkable changes. C = normal chondrocytes with normal periphery staining, spatial distribution and background staining intensity [H & E Stain x 400].

## DISCUSSION

In this study, the marked changes observed in the physical appearances of the rats in the model group is an important indication of RA in the research animals, which was also assessed and monitored through the notable progression of symmetrical swelling of multiple joints, limited mobility, fur discoloration and weight loss of research animals. The skin ulceration and scab formation identified was not an indication of RA, however it can be attributed to have come from possible errors in the animal handling technique adopted. The disparity in the body weight of rats in the normal control group and model group at day 16 can be attributed to the onset of RA disease.[18] After treatment was initiated for 12 weeks, the low dose group B and positive control group A showed no remarkable weight gain while group C and D had appreciable weight gain. The unremarkable weight gain exhibited by group B could be as a result of the extract dosage, due to the difference seen in groups C and D. Again, group D had better results when compared to group C and group B, thus reiterating the stance of the effect of high extract dosage. This suggests potential synergistic effect of the *Jatropha tanjorensis* plant extract in achieving sustained clinical remission in rheumatoid arthritis [19]. It could be attributed to the antioxidant properties of *J. tanjorensis* or its ameliorative effect on gastrointestinal disorders caused by NSAIDs [2, 10].

The anterior-posterior and lateral diameters of the ankles measured served as an analytical tool to assess the treatment phase and administration of the ethanolic *J. tanjorensis* leaf extract [18]. The difference noted on day 16 between the normal control group and model group was a clear indication of onset of RA, however on day 93 long after treatment commenced, group A and B exhibited results suggestive of poor response to the treatment. The observed effective response to treatment was more pronounced in groups C and D, indicating a better treatment response to higher dosage of extract. Group C (medium dose) responded but was not as efficient as the response observed in group D (high dose).

The results from the Modified Mankin scoring system analysis of the micrograph on *J. tanjorensis* extract effect on rheumatoid arthritis suggest that the extract may not have synergistic effect at low dose levels. This was evident in the marked level of cartilage destruction demonstrated by superficial fibrillation of chondrocytes and clustering, with increased intensity of periphery and background staining of the cartilages and matrix respectively in the low dose (0.5g/kg) treatment group. However, increasing the extract dose level to 1.0g/kg improved the features observed in the cartilage, and instead, apart from superficial fibrillation of the chondrocytes, it demonstrated diffuse hyper-cellular spatial arrangement of the chondrocytes with slightly enhanced chondrocyte periphery and matrix background staining. The improved features observed in the cartilages the of joints of rats treated with increased doses of *J. tanjorensis* leaf extracts suggests that the extract could cause the stimulation of the activities of the Ibuprofen and Sulfasalazine earlier than normal due to the anti-inflammatory and antioxidant activities which suppresses oxidative reactions [20]. However, the ameliorative effect of the leaf extracts on rheumatoid arthritis could be attributed to constituent phytochemicals such as terpenoids, saponins or flavonoids and tannins with known antioxidant and anti-inflammatory properties [13, 14, 21].

## CONCLUSION

In conclusion, Rheumatoid Arthritis has been established to be one of the leading causes of debility and morbidity in the world. The treatment modalities are limited to conventional drugs (CD) such as disease-modifying anti-rheumatic drugs (DMARDs) and non-steroidal anti-inflammatory drugs (NSAIDs) as newer medications such as biologics are not easily affordable due to lack of purchasing power and poverty amongst patients. Complementary and alternative medicine seem to be the order of the day in people suffering from chronic pain, as in RA; which includes traditional and herbal medications. This growing interest in alternative medicine clearly indicates a need for research to go beyond identifying herbs but as well searching for a better and effective anti-rheumatic and anti-inflammatory drugs from plants. There is equally a very important need to standardize dosages of newer herbs identified to be therapeutic. It is also recommended that the side effects of such herbs be finely elucidated especially with regards their effects on certain organs in the body such as the kidneys and liver for obvious reasons.

## ETHICS STATEMENT

Patient consent for publication Not applicable

### Ethical approval

Animal experiment protocol was approved by the Research and Ethical Clearance Committee, Animal Research Ethics Committee, Faculty of Basic Medical Sciences, College of Medicine, Enugu State University of Science and Technology (<u>ESUCOM/FBMS/ETR/2021/007).</u>

## COMPETITING INTEREST

None declared.

## ACKNOWLEDGEMENT

Animal house and care was provided by Enugu State University College of Medicine.

